# Suitability of different mapping algorithms for genome-wide polymorphism scans with Pool-Seq data

**DOI:** 10.1101/052845

**Authors:** Robert Kofler, Anna Maria Langmüller, Pierre Nouhaud, Kathrin Anna Otte, Christian Schlöetterer

## Abstract

The cost-effectiveness of sequencing pools of individuals (Pool-Seq) provides the basis for the popularity and wide-spread use of this method for many research questions, ranging from unravelling the genetic basis of complex traits to the clonal evolution of cancer cells. Because the accuracy of Pool-Seq could be affected by many potential sources of error, several studies determined, for example, the influence of the sequencing technology, the library preparation protocol, and mapping parameters. Nevertheless, the impact of the mapping tools has not yet been evaluated. Using simulated and real Pool-Seq data, we demonstrate a substantial impact of the mapping tools leading to characteristic false positives in genome-wide scans. The problem of false positives was particularly pronounced when data with different read lengths and insert sizes were compared. Out of 14 evaluated algorithms novoalign, bwa mem and clc4 are most suitable for mapping Pool-Seq data. Nevertheless, no single algorithm is sufficient for avoiding all false positives. We show that the intersection of the results of two mapping algorithms provides a simple, yet effective strategy to eliminate false positives. We propose that the implementation of a consistent Pool-seq bioinformatics pipeline building on the recommendations of this study can substantially increase the reliability of Pool-Seq results, in particular when libraries generated with different protocols are being compared.

## Introduction

Sequencing pools of individuals (Pool-Seq) is a cost efficient approach for generating genome-wide polymorphism data, which is enjoying increasing popularity [reviewed in Schlötterer et al. (2014)]. Pool-Seq was for example used to unravel the genetic basis of complex traits (Bastide et al., 2013; Cheeseman et al., 2015), identify loci contributing to local adaptation (Lamichhaney et al., 2012; Turner et al., 2010), trace beneficial loci during experimental evolution (Lang et al., 2013; Orozco-terWengel et al., 2012; Tobler et al., 2013), identify positively selected loci in populations (Bergland et al., 2014; Kofler et al., 2012; Nolte and Schlötterer, 2008), find genes selected during domestication (Axelsson et al., 2013; Rubin et al., 2010), study the invasion of transposable elements (Kofler et al., 2015a), investigate clonal evolution in cancer (Ding et al., 2012) and to identify causative mutations in forward genetic screens (Schneeberger et al., 2009). With this rapid gain in popularity it is important to ensure a reliable analysis of Pool-Seq data. Several studies investigated various aspects that potentially affect the accuracy of Pool-Seq, including the sequencing platform (Rellstab et al., 2013), the reference genome (Nevado et al., 2014), the parameters used for aligning the reads (Kofler et al., 2011a), the sequencing depth (Ferretti et al., 2013; Kofler and Schlötterer, 2014), the pool size (Futschik and Schlötterer, 2010; Gautier et al., 2013) and the library preparation protocol (Kofler et al., 2015b).

However, until now the impact of the mapping algorithm used for aligning Pool-Seq data has not been studied in sufficient detail. Here, we show that the mapping algorithm can have a profound effect leading to erroneous signals of allele frequency differences between libraries. We systematically compared the performance of 14 different alignment algorithms using both simulated and real Pool-Seq data. Of the tested algorithms clc4, novoalign and bwa mem consistently produced the most reliable results with Pool-Seq data. Nevertheless, no single alignment algorithm avoids all artefacts, but by intersecting the results of two alignment tools, the vast majority of artifactual outliers can be avoided.

## 1 Results

Genome wide polymorphism scans with Pool-Seq data are becoming increasingly used in population genomic research. Typically, these studies use genome-wide Pool-Seq data to identify marked outlier loci in pairwise comparisons between population samples. For example, loci contributing to local adaptation are identified by significantly different allele frequencies between populations (Lamichhaney et al., 2012; Turner et al., 2010). This focus on outlier loci makes genome-wide scans susceptible to technical problems that could generate outlier artefacts. We found that the mapping algorithms for aligning Pool-Seq data may be an important source of outlier artefacts (fig. 1). Comparing allele frequencies between two Pool-Seq libraries prepared from identical genomic DNA, but with different insert size and read length, we found a substantial number of outlier loci, despite no differences between the libraries are expected (fig. 1A,B).

**Figure 1:**
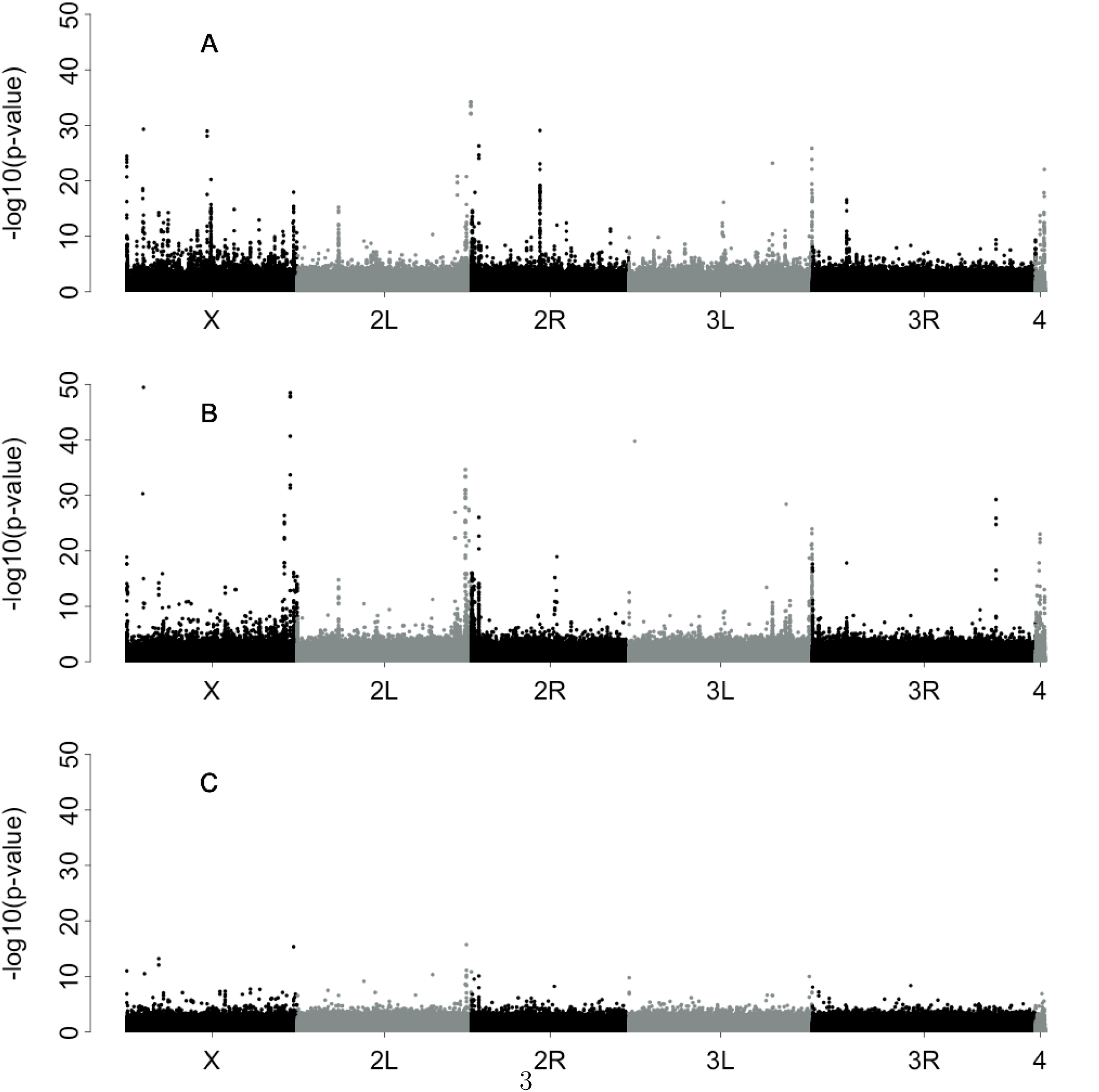
Manhattan plots indicating the significance of allele frequency differences between Pool-Seq libraries when the same genomic DNA is sequenced. Two Illumina paired-end sequencing libraries with different read length and insert sizes were prepared from a pool of 250 *D. simulans* individuals. Reads were mapped to the reference genome and the significance of differences in allele frequencies between the two libraries were computed (Fisher’s exact test). Despite no significant allele frequency differences are expected we found pronounced outlier peaks using bwa aln (A) or novoalign(g) (B) for mapping the reads. Importantly, outlier peaks found with these two alignment algorithms are at different genomic sites. Hence, intersecting the results of these two algorithms by plotting the lowest obtained p-value at each site removes the vast majority of outlier peaks (C).

To overcome this problem we set out to identify alignment algorithms that are most suitable for genome-wide outlier scans using Pool-Seq data. We tested seven semi-global alignment algorithms, where the entire read is required to match [bowtie2(g), bwa aln, clc4(g), mrfast, ngm(g), novoalign(g), segemehl], and seven local alignment algorithms, where only a part of the read needs to match [bwa sw, bwa mem, clc4(l), gsnap, ngm(l), novoalign(l); for an overview see table 6]. (Alkan et al., 2010; CLC bio, 2015; Hoffmann et al., 2009; Langmead and Salzberg, 2012; Li and Durbin, 2009, 2010; Novocraft, 2014; Sedlazeck et al., 2013; Wu and Nacu, 2010). With several tools, like ngm or bowtie2, supporting both semi-global and local alignments, we indicate the pertinent algorithm in brackets [e.g.: ngm(g): semi-global alignment, ngm(l): local alignment].

We first tested the overall performance of the alignment algorithm using simulated data sets. We generated template sequences with SNPs and indels at known positions and then simulated uniformly distributed paired ends from these templates such that true SNPs are spaced exactly 100bp and have a population frequency of 0.5 (fig. 2). Note that indels are in linkage disequilibrium with SNPs to identify biased allele frequency estimates resulting from mapping of reads with indels.

**Figure 2:**
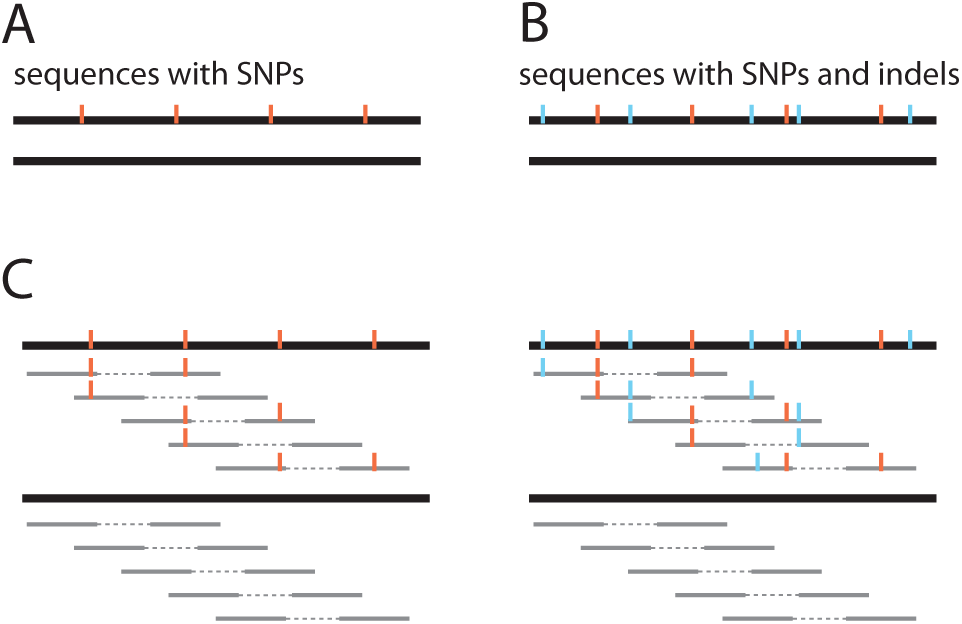
Overview of simulated Pool-Seq data sets. Based on a 2 Mbp region of *D. melanogaster* chromosome 2R, we simulated a pair of sequences with one sequence having a SNP (red) every 100bp (A) and a pair of sequences with one sequence having, in addition to the SNPs, an indel (blue) with random position and length between adjacent SNPs (B). Using these sequences as templates we simulated uniformly distributed paired ends (grey; C) resulting in SNPs with known positions and frequency (0.5).

We evaluated the mapping algorithms with three different paired end data sets: i) a data set representing optimal conditions (2x100bp paired ends; insert size 100±0bp; error rate of 0%; no indels; fig. 2A), ii) a data set with indels and variation of the distance between paired ends (2x100bp paired ends; insert size 100 ± 40bp; error rate of 0%; indels; fig. 2B) and a dataset with indels and a high error rate (polymorphism) of 5% (2x100bp paired ends with an insert size of 100±0bp; error rate of 5%; indels; fig. 2B). For all data sets a coverage of 200 per site was targeted (≈2 million paired ends per data set). We evaluated the performance of the mapping algorithms based on four criteria: the number of true positive SNPs, the number of false positive SNPs, the average frequency of the reference allele (should be 0.5) and the number of extreme outlier loci with highly inaccurate allele frequency estimates (ƒ ≥ 0.9 or ƒ ≤ 0.1). These criteria could, for example, be important in genome-wide scans to detect signatures of selection with Pool-Seq data, where a reliable identification of regions having low polymorphism, a hallmark of positive selection, requires precise identification of the SNPs and accurate estimates of allele frequencies.

We compared the performance of the mapping algorithms with and without filtering for quality criteria, such as paired end reads and mapping quality [≥ 20; a low mapping quality suggest that the read is ambiguously mapped (Li et al., 2008)] and found that filtering consistently leads to reduced numbers of false positive SNPs and more accurate allele frequency estimates (supplementary table 1). This observation is in agreement with previous work showing that quality filtering can reduce the number of false positive SNPs (Li et al., 2008). We note, however, that quality filtering also leads to fewer true positive SNPs (supplementary table 1).

Quality filtering also affected the coverage distribution. Fewer sites had a higher coverage than simulated in filtered data (supplementary figure 1), which is likely due to smaller numbers of ambiguously mapped reads that stochastically accumulate in some genomic regions. For mrfast quality filtering resulted in a severe shift of the coverage distribution, halving the average coverage (supplementary figure 1). The distribution of mapping qualities differed between mapping algorithms (supplementary fig. 2) which is likely due to distinct algorithms for computing mapping qualities. Since the accuracy of allele frequency estimates was substantially better for filtered data sets, we rely on quality filtered reads for the remaining manuscript. Summarizing the results for all three simulated data sets we found that clc4(g), novoalign(g), bwa bwasw, clc4(l), gsnap and novoalign(l) showed the best performance while bowtie2(g), mrfast, ngm(g) and segemehl showed the worst (table 1; for results with unfiltered data see supplementary table 2). The average reference allele frequency of most alignment algorithms was above 0.5 indicating a bias towards the reference allele [see also Degner et al. (2009); Kofler et al. (2011a)]. After quality filtering mrfast had a substantial bias against the reference allele (supplementary table 2).

**Table 1:** Suitability of mapping algorithms for performing genome wide polymorphism scans with Pool-Seq data. Ideally, a mapping algorithm should enable to identify all true positive SNPs (TP; 19.999 were simulated) and to estimate the allele frequencies accurately (*μ*_ƒ_ average frequency of the reference allele; all SNPs were simulated with ƒ = 0.5) while avoiding the identification of false positive SNPs (FP) and extreme outlier SNPs, with highly inaccurate allele frequency estimates (OL; ƒ > 0.9 or ƒ < 0.1). We tested the algorithm with three different data sets. For each benchmark the three best performing (green) and the three worst performing (red) algorithm were marked. The overall suitability of each mapping algorithm was determined (count top - count worst; top ≥ 2; worst ≤ −2) and algorithms were marked accordingly. False positive SNPs were not used as benchmark for the data with the high error rate. best case: 2x100bp paired ends with an insert size of 100±0bp, indel - insert size: 2x100bp paired ends with an insert size of 100±40bp and indels between the SNPs, indel - error rate: 2x100bp paired ends with an insert size of 100±0bp, indels between the SNPs and an error rate of 5%.

| algorithm | best case |  |  |  | indel - insert size |  |  |  | indel - error rate |  |  |  |
| --- | --- | --- | --- | --- | --- | --- | --- | --- | --- | --- | --- | --- |
| | TP | FP | $\mu_f$ | OL | TP | FP | $\mu_f$ | OL | TP | FP | $\mu_f$ | OL |
| bowtie2(g) | 12649 | 0 | 0.556 | 367 | 12561 | 23222 | 0.576 | 642 | 10461 | 423k | 0.855 | 2940 |
| bwa aln | 15822 | 5 | 0.501 | 1 | 15978 | 36634 | 0.538 | 184 | 14980 | 487k | 0.583 | 189 |
| clc4(g) | 16667 | 135 | 0.504 | 2 | 16787 | 15376 | 0.511 | 155 | 16655 | 1527k | 0.504 | 142 |
| mrfast | 1277 | 256 | 0.083 | 1109 | 1852 | 7038 | 0.120 | 1510 | 14920 | 661k | 0.778 | 1032 |
| ngm(g) | 10536 | 28 | 0.495 | 10 | 10337 | 8290 | 0.529 | 134 | 9735 | 802k | 0.521 | 87 |
| novoalign(g) | 16508 | 20 | 0.501 | 1 | 16630 | 10457 | 0.508 | 124 | 16482 | 1506k | 0.507 | 148 |
| segemehl | 19984 | 62k | 0.605 | 1214 | 19880 | 96448 | 0.610 | 1367 | 19987 | 1957k | 0.640 | 1304 |
| bowtie2(l) | 11078 | 0 | 0.597 | 599 | 11290 | 281 | 0.616 | 846 | 11193 | 855k | 0.534 | 97 |
| bwa bwsw | 14099 | 0 | 0.524 | 2 | 14259 | 1134 | 0.583 | 73 | 13807 | 1101k | 0.645 | 219 |
| bwa mem | 11418 | 0 | 0.502 | 7 | 16558 | 10177 | 0.509 | 117 | 14451 | 1313k | 0.349 | 3179 |
| clc4(l) | 16642 | 62 | 0.513 | 6 | 16730 | 2388 | 0.519 | 96 | 16606 | 1514k | 0.516 | 155 |
| gsnap | 16621 | 250 | 0.520 | 5 | 17034 | 6607 | 0.526 | 65 | 17267 | 1491k | 0.548 | 221 |
| ngm(l) | 10458 | 0 | 0.504 | 13 | 10186 | 1208 | 0.537 | 95 | 9610 | 775k | 0.526 | 85 |
| novoalign(l) | 16446 | 19 | 0.523 | 3 | 16504 | 361 | 0.526 | 18 | 16396 | 1483k | 0.527 | 168 |

Next we compared allele frequency estimates between samples, an approach that is typically used to identify loci responsible for local adaption. We investigated the sensitivity of the alignment algorithm to i) differences of the inner distance between paired ends (inner distances 100 ± 20bp vs. 300 ± 60bp) ii) differences in read length (read length 100bp vs. 50bp) and iii) differences in the error rates (error rates 1% vs. 5%) (table 1). Uniformly distributed paired ends were simulated from the template sequences having SNPs and indels (fig. 2B). Allele frequency differences between samples were measured using *F*_ST_. Values of *F*_ST_ range from 0 to 1, where 0 indicates no differentiation between samples (populations) and 1 indicates complete differentiation (fixation for alternative alleles) (Hartl and Clark, 1997). As all paired ends have a uniform genomic distribution and were derived from the same template sequences only small allele frequency differences are expected between sam-ples. A perfect alignment algorithms would detect all positive SNPs (*TP* = 19.999) and yield a low *F*_ST_ for all SNPs (*F*_ST_ = 0). Based on the simulated data clc4(g), novoalign(g), bwa mem, clc4(l) and novoalign(l) showed the best performance whereas mrfast, ngm(g), bowtie2(l) and bwa bwasw performed worst (table 1; for allele frequency differences with false positive SNPs see supplementary table 3). We noted substantial allele frequency differences when the same data were mapped as paired-end and as single-end reads and then compared against each other (supplementary table 4). ngm(g) and ngm(l) were most suitable for such comparisons between paired and single end reads (supplementary table 4).

**Table 2:** Comparison of allele frequency differences between simulated Pool-Seq data sets with different mapping algorithms. We simulated different paired end Pool-Seq libraries, mapped the reads and compared the allele frequencies between the libraries using *F*_ST_. With this procedure we evaluated the sensitivity of the alignment algorithm to differences in the distance between paired ends (id), differences in the read length (rl) and differences in the error rates (e). As all libraries were derived from identical template sequences (templates with SNPs and indels) no significant allele frequency differences were expected (*F*_ST_ = 0). We estimated the number of true positive SNPs for which allele frequencies could be compared (TP) and the lowest *F*_ST_-values in the 0.1% and 10% quantiles with the most differentiated SNPs. For each benchmark we highlighted the three best (green) and three worst (red) performing algorithm. The overall suitability of each mapping algorithm was determined (count top - count worst; top ≥ 3; worst ≤ −3) and algorithms were marked accordingly. id100, rl100, e1%: 2x100bp paired ends, insert size 100±20bp, error rate 1%; id300: 2x100bp paired ends, insert size 300 ± 60bp, error rate 1%; rl50: 2x50bp paired ends, insert size 100 ± 20bp, error rate 1%; e5%: 2x100bp paired ends, insert size 100 ± 20bp, error rate 5%

| algorithm | id100 vs. id300 |  |  | rl100 vs. rl50 |  |  | e1% vs. e5% |  |  |
| --- | --- | --- | --- | --- | --- | --- | --- | --- | --- |
|  | TP | 10% | 0.1% | TP | 10% | 0.1% | TP | 10% | 0.1% |
| bowtie2(g) | 12468 | 0.021 | 0.358 | 12474 | 0.027 | 0.351 | 11778 | 0.251 | 0.476 |
| bwa aln | 14128 | 0.008 | 0.334 | 14106 | 0.014 | 0.279 | 14860 | 0.078 | 0.486 |
| clc4(g) | 12415 | 0.006 | 0.286 | 11067 | 0.008 | 0.201 | 12664 | 0.004 | 0.115 |
| mrfast | 10752 | 0.019 | 0.46 | 9712 | 0.055 | 0.500 | 11704 | 0.212 | 0.603 |
| ngm(g) | 8463 | 0.006 | 0.143 | 7122 | 0.030 | 0.468 | 7878 | 0.017 | 0.175 |
| novoalign(g) | 16093 | 0.004 | 0.289 | 15263 | 0.007 | 0.232 | 16415 | 0.003 | 0.081 |
| segemehl | 14097 | 0.005 | 0.223 | 13394 | 0.010 | 0.216 | 11550 | 0.011 | 0.136 |
| bowtie2(l) | 11158 | 0.031 | 0.359 | 11280 | 0.039 | 0.301 | 10663 | 0.024 | 0.668 |
| bwa bwasw | 13694 | 0.016 | 0.363 | 11433 | 0.262 | 0.456 | 13661 | 0.020 | 0.215 |
| bwa mem | 15990 | 0.005 | 0.290 | 15104 | 0.007 | 0.156 | 16288 | 0.004 | 0.075 |
| clc4(l) | 12415 | 0.006 | 0.286 | 11067 | 0.008 | 0.201 | 12664 | 0.004 | 0.115 |
| gsnap | 16363 | 0.007 | 0.231 | 12350 | 0.041 | 0.385 | 13615 | 0.044 | 0.372 |
| ngm(l) | 9665 | 0.006 | 0.142 | 9110 | 0.026 | 0.453 | 9079 | 0.018 | 0.167 |
| novoalign(l) | 16057 | 0.005 | 0.279 | 15191 | 0.009 | 0.243 | 16381 | 0.003 | 0.081 |

Simulated data may not capture all the properties of real data such as reads having different lengths (after trimming), variable base qualities along reads and biases in sequencing errors. Therefore we also evaluated the performance of different alignment algorithms based on *F*_ST_ between samples using real data.

We used two libraries with different read length and insert size prepared from the same genomic DNA (library 1: 2x76bp paired ends, median insert size = 232bp; library 2: 2x120bp paired ends, median insert size = 396; both prepared from pooled *D. simulans* flies; see Material and Methods), trimmed low quality regions from the 3’-ends of reads and compared allele frequency differences between the samples using *F*_ST_. As both libraries were prepared from the same genomic DNA only small allele frequency differences were expected between the samples (*F*_ST_ = 0). Novoalign(g), bwa mem, and novoalign(l) showed the best performance while clc(g), mrfast, segemehl, gsnap and ngm(l) performed worst (table 3).

**Table 3:** Comparison of allele frequency differences between real Pool-Seq data sets with different mapping algorithms. We compared allele frequencies between two paired end libraries with different read length and insert size that were prepared from the same genomic DNA (pooled *D. simulans* flies). We determined the number of SNPs for which allele frequencies could be compared (*c*) and the lowest *F*_ST_-values in different quantiles with the most differentiated SNPs. For each benchmark the three top (green) and three worst (red) performing algorithms are highlighted. The overall suitability of each mapping algorithm was determined (count top - count worst; top ≥ 2; worst ≤ −2) and algorithms were marked accordingly. The number of SNPs (c) was not used as a benchmark as the true SNPs are not known. mrfast generated an invalid output file with these data (an uniform read length was reported despite these reads having varying read lengths).

| algorithm | c | 10% | 1% | 0.10% | 0.01% | 0.001% |
| --- | --- | --- | --- | --- | --- | --- |
| bowtie2(g) | 4914k | 0.074 | 0.154 | 0.274 | 0.429 | 0.651 |
| bwa aln | 5030k | 0.064 | 0.137 | 0.253 | 0.437 | 0.660 |
| clc4(g) | 6610k | 0.064 | 0.139 | 0.265 | 0.464 | 0.692 |
| mrfast | na | na | na | na | na | na |
| ngm(g) | 8427k | 0.068 | 0.142 | 0.263 | 0.447 | 0.685 |
| novoalign(g) | 4964k | 0.057 | 0.125 | 0.228 | 0.373 | 0.578 |
| segemehl | 4866k | 0.072 | 0.151 | 0.285 | 0.492 | 0.750 |
| bowtie2(l) | 4745k | 0.062 | 0.133 | 0.247 | 0.419 | 0.608 |
| bwa bwsw | 4531k | 0.068 | 0.148 | 0.266 | 0.415 | 0.590 |
| bwa mem | 4786k | 0.061 | 0.132 | 0.248 | 0.409 | 0.607 |
| clc4(l) | 4714k | 0.064 | 0.143 | 0.275 | 0.464 | 0.670 |
| gsnap | 4897k | 0.068 | 0.155 | 0.303 | 0.501 | 0.693 |
| ngm(l) | 5050k | 0.064 | 0.143 | 0.283 | 0.492 | 0.708 |
| novoalign(l) | 4387k | 0.060 | 0.132 | 0.238 | 0.387 | 0.570 |

In summary, when comparing the results of the previous evaluations, we conclude that clc4(g), novoalign(g), bwa mem, clc4(l) and novoalign(l) are the most suitable alignment algorithm for Pool-Seq data whereas bowtie2(g), mrfast, ngm(g), segemehl, bowtie2(l), ngm(l) did not perform as well (table 4).

**Table 4:**
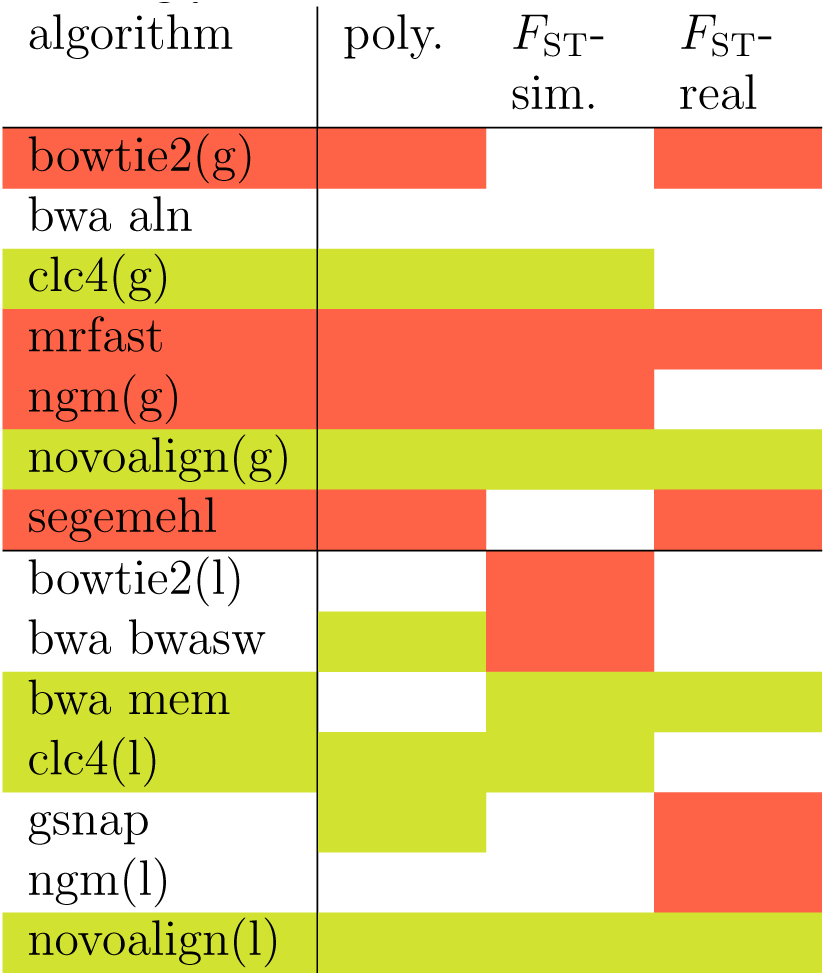
Comparision of alignment algorithms for Pool-Seq data: summary across data sets. Tables shows an overview of the results of the previous evaluations: overall suitability (poly: table 1), allele frequency differences using simulated data (*F*_ST_-sim.: table 1) and allele frequency differences using real data (*F*_ST_-real: table 3). The overall suitability of each mapping algorithm was determined (count top - count worst; top ≥ 2; worst ≤ −2) and algorithms were marked accordingly.

Despite novoalign(g) being one of the most suitable algorithms for Pool-Seq data, a substantial number of artifactual outlier peaks can still be found when comparing the allele frequency between the *D. simulans* libraries (fig. 1). The comparison of different mappers indicated that outlier artefacts are frequently specific to the alignment algorithm (Fig. 1; supplementary fig. 3, 4). We reasoned therefore that an intersection of two mappers, recording for every SNP only the least significant result found by any mapper, could overcome this problem. Intersecting the results of bwa and novoalign (fig. 1A,B), the number of outlier peaks could be substantially reduced (fig. 1C). We also tested whether intersecting the results of different mappers preserves the targets of selection using data from an experimental evolution study for C-virus resistance in *D. melanogaster* (Martins et al., 2014) and found that the most differentiated loci identified by Martins et al. (2014) were retained (supplementary fig. 5). Hence, intersecting the results of different mappers is a viable strategy for minimizing the number of artefacts while preserving the targets of selection. To identify the most suitable combination of mapping algorithms we used the data from the pooled *D. simulans* flies, computed all pairwise intersections of the algorithms and benchmarked them using the number of SNPs and the 0.001% quantile of most differentiated SNPs (table 5). ngm(l) combined with bowtie2(g) yielded the least pronounced outlier peaks with about 4.15 million shared SNPs [table 5; for Manhattan plots see supplementary fig. 6]. We note, however, that the best combination of alignment algorithms depends on the threshold–with the 0.01% quantile novoalign(l) and bowtie2(g) are the best combination (supplementary table 5; supplementary fig. 8). Interestingly, combining the two algorithm that were individually the most suitable for Pool-Seq data, novoalign(g) and bwa mem (table 4), did not lead to a marked reduction of outlier peaks (table 5); supplementary fig. 7). We hypothesize that this could be due to a high similarity of the alignment algorithms.

**Table 5:**
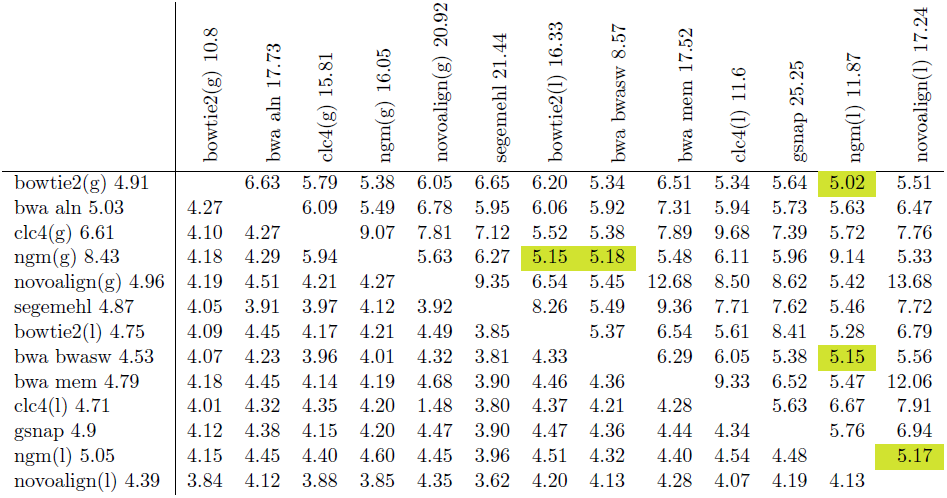
Reduction of mapping artefacts by the intersection of mapping algorithms. Two Illu-mina paired end data sets with different insert sizes and read length were derived from pooled genomic DNA (natural *D. simulans* population) and mapped to the reference genome. Allele frequency differences between the libraries were computed using Fisher’s exact test [-log(p-value) = ƒ*et*-*value*]. To test which combination of alignment algorithms most efficiently reduces outlier peaks we intersected all pairwise combinations of alignment algorithms, i.e. we use SNPs identified with both algorithms and use the lowest ƒ*et*-*value* found with any of the two algorithms. Below the diagonal we report the number of SNPs (in million) common to both algorithms. For comparision, the number of SNPs identified with a single mapping algorithm are shown next to the list of mappers on the left side. Above the diagonal we report the lowest ƒ*et*-*value* among the 0.001% most differentiated SNPs (around 40 SNPs). For comparision, the corresponding ƒ*et*-*values* obtained with a single mapping algorithm are shown next to the list of mappers on the upper side. We marked the five best combinations yielding the least pronounced outlier loci (green).

|  | bowtie2(g) 10.8 | bwa aln 17.73 | clc4(g) 15.81 | ngm(g) 16.05 | novoalign(g) 20.92 | segemehl 21.44 | bowtie2(l) 16.33 | bwa bwasm 8.57 | bwa mem 17.52 | clc4(l) 11.6 | gsnap 25.25 | ngm(l) 11.87 | novoalign(l) 17.24 |
| --- | --- | --- | --- | --- | --- | --- | --- | --- | --- | --- | --- | --- | --- |
| bowtie2(g) 4.91 |  | 6.63 | 5.79 | 5.38 | 6.05 | 6.65 | 6.20 | 5.34 | 6.51 | 5.34 | 5.64 | 5.02 | 5.51 |
| bwa aln 5.03 | 4.27 |  | 6.09 | 5.49 | 6.78 | 5.95 | 6.06 | 5.92 | 7.31 | 5.94 | 5.73 | 5.63 | 6.47 |
| clc4(g) 6.61 | 4.10 | 4.27 |  | 9.07 | 7.81 | 7.12 | 5.52 | 5.38 | 7.89 | 9.68 | 7.39 | 5.72 | 7.76 |
| ngm(g) 8.43 | 4.18 | 4.29 | 5.94 |  | 5.63 | 6.27 | 5.15 | 5.18 | 5.48 | 6.11 | 5.96 | 9.14 | 5.33 |
| novoalign(g) 4.96 | 4.19 | 4.51 | 4.21 | 4.27 |  | 9.35 | 6.54 | 5.45 | 12.68 | 8.50 | 8.62 | 5.42 | 13.68 |
| segemehl 4.87 | 4.05 | 3.91 | 3.97 | 4.12 | 3.92 |  | 8.26 | 5.49 | 9.36 | 7.71 | 7.62 | 5.46 | 7.72 |
| bowtie2(l) 4.75 | 4.09 | 4.45 | 4.17 | 4.21 | 4.49 | 3.85 |  | 5.37 | 6.54 | 5.61 | 8.41 | 5.28 | 6.79 |
| bwa bwasm 4.53 | 4.07 | 4.23 | 3.96 | 4.01 | 4.32 | 3.81 | 4.33 |  | 6.29 | 6.05 | 5.38 | 5.15 | 5.56 |
| bwa mem 4.79 | 4.18 | 4.45 | 4.14 | 4.19 | 4.68 | 3.90 | 4.46 | 4.36 |  | 9.33 | 6.52 | 5.47 | 12.06 |
| clc4(l) 4.71 | 4.01 | 4.32 | 4.35 | 4.20 | 1.48 | 3.80 | 4.37 | 4.21 | 4.28 |  | 5.63 | 6.67 | 7.91 |
| gsnap 4.9 | 4.12 | 4.38 | 4.15 | 4.20 | 4.47 | 3.90 | 4.47 | 4.36 | 4.44 | 4.34 |  | 5.76 | 6.94 |
| ngm(l) 5.05 | 4.15 | 4.45 | 4.40 | 4.60 | 4.45 | 3.96 | 4.51 | 4.32 | 4.40 | 4.54 | 4.48 |  | 5.17 |
| novoalign(l) 4.39 | 3.84 | 4.12 | 3.88 | 3.85 | 4.35 | 3.62 | 4.20 | 4.13 | 4.28 | 4.07 | 4.19 | 4.13 |  |

## 2 Discussion

Here, we performed a comprehensive analysis of different alignment algorithms for Pool-Seq data. The evaluation of alignment algorithms is complicated by several issues. First, the mapping quality is computed differently between algorithms (supplementary fig. 2). Thus, the fraction of reads filtered by requiring a certain minimum quality (we used 20) varies among the alignment tools. The fraction of filtered reads will affect both the number of identified true positive SNPs and the accuracy of the allele frequency estimates: more mapped reads result in a higher number of true SNPs but also the number of ambiguously mapped reads is increased, which distorts the allele frequency estimates. The tradeoff between optimizing the recovery of true SNPs and accuracy of the allele frequency estimates is particularly pronounced for segemehl: no reads could be quality filtered since all reads have a mapping quality of 255, resulting in the highest number of true positive SNPs but poor allele frequency estimates (table 1). Despite this complication, we considered quality filtering of reads essential as this substantially improves allele frequency estimates from Pool-Seq data (for unfiltered results see supplementary table 2). Interestingly, the best performing algorithm (e.g. novoalign and clc) identified the highest number of true positive SNPs and yielded the most accurate allele frequency estimates (table 1), which suggests that the superior performance of these tools is robust with respect to the tradeoff introduced by quality filtering.

The choice of alignment parameters is a challenge for the comparison of different mapping algorithms. Whenever feasible, we used default parameters and only modified them when we considered it necessary to ensure an unbiased comparision (e.g. when the error rate exceeded the number of allowed mismatches or when the insert size was larger than the maximum insert size; see Material and Methods). We note, however, that the performance of each of these algorithms may be improved by fine-tuning the parameters. For example, the performance of bwa aln was substantially improved by using parameters optimized for Pool-Seq data Kofler et al. (2011a) (supplementary table 6). While the optimization of mapping parameters for all 14 alogrithms is clearly beyond the scope of this manuscript, we made all data, including the simulated ones, publicly available to allow testing the performance of different mappers and parameters with these data sets.

Out of the 14 algorithms tested clc4(g), novoalign(g), bwa mem, clc4(l) and novoalign(l) are the most suitable ones for Pool-Seq data. The superior performance of novoalign is in agreement with previous work which found that novoalign yields highly accurate alignments and SNP calls (Bao et al., 2014; Li and Homer, 2010; Nielsen et al., 2011).

The most striking influence of different alignment algorithms was noted for experimental data differing in insert size and read length. Comparing different libraries from the same genomic DNA, we identified substantial outliers, some of them clustering in peaks which indicate allele frequency differences at multiple neighbouring sites. Since such peaks are a typical signal in genome-wide outlier scans, such as Pool-GWAS or E&R studies, these artefacts may lead to false conclusions. Similar artefacts were also seen when the data were mapped as single reads (supplementary fig. 9), suggesting that this is not an artefact of paired end mapping. Assuming that true allele frequency differences between samples should be identified with most alignment tools, whereas artefacts should only be found with a few algorithms, we propose intersecting multiple alignment algorithms. We noticed a clear improvement when intersecting two alignment algorithms but, depending on the evaluation criteria, different pairs of algorithms perform best. These results are consistent with other studies, which also found that the combination of mapping algorithms and/or variant calling pipelines may yield superior results (Bao et al., 2014; Field et al., 2015; O’Rawe et al., 2013).

Our approach to intersect algorithms is based on the least significant allele frequency differences between two samples. It is straight forward to extend this approach to studies that rely on multiple samples, such as replicated Pool-GWAS experiments or E&R studies (for example see supplementary fig. 5), provided that it is feasible to collapse allele frequency differences between multiple samples into a single representative measure [e.g. p-value from a cmh-test (Orozco-terWengel et al., 2012)]. In this case again the least significant value found by any mapper may be used. However, this strategy cannot be applied to Pool-Seq data from single populations (e.g. Asgharian et al., 2015; Boitard et al., 2013; Nolte et al., 2012). One possibility to avoid mapping artefacts for single population Pool-Seq data may be to filter SNPs with incongruent allele frequency estimates among multiple mappers. Given that most artefacts were observed when libraries with different insert sizes and read lengths were compared (fig. 2 vs. supplementary fig. 10), we recommend using a single consistent sequencing strategy for all Pool-Seq libraries, when ever possible. We additionally propose to use a single consistent mapping pipeline for all Pool-Seq data, as mixing samples aligned with different tools, algorithms, parameters or even versions of the same tool, leads to elevated levels of outlier peaks (supplementary table 6).

## 3 Material and Methods

### 3.1 Alignment algorithms

We tested seven semi-global alignment algorithms, where the entire read is required to match, and seven local alignment algorithms, where only a part of the read needs to match (table 6). For tools that support semi-global as well as local alignments we evaluated the suitability of both algorithms (table 6). We also included gsnap (Wu and Nacu, 2010) into our study, despite this tool was designed for aligning RNA-Seq data (i.e. alignments with large gaps to allow for spliced introns). We also aimed to include gem (Marco-Sola et al., 2012), batalign (Lim et al., 2015), stampy (Lunter and Goodson, 2011) and soap2 (Li et al., 2009b) into our study but were not able to run these tools on our computational infrastructure (Mac Pro; batalign: did not respond, gem: compilation failed, stampy: compilation failed due to missing files, soap2: segmentation fault while indexing the reference genome). If possible we used default parameters for all tools and only deviated from these settings when deemed necessary to ensure an unbiased comparison of the alignment algorithms (table 6). With Bowtie2 we set the maximum fragment length of paired ends (-X) to 1500. For bwa we used version 0.7.4 for the mem and bwasw algorithm and version 0.6.2 for the aln algorithm. This was necessary as bwa aln 0.7.4 reports a segmentation fault when aligning some data sets (e.g. the *D. simulans* libraries) whereas the mem algorithm was not available for bwa version 0.6.2. For clc4 we interleaved the sequences of the two fastq files (-i), activated the paired end mode (-p), set the orientation of the paired ends to forward followed by backward (fb) and measured the distance between paired ends from start-to-start (ss). As the performance of clc4 is highly sensitive to the provided minimum distance (min) and maximum distance (max) between paired ends we provided the most suitable setting for each alignment (simulated data, read length 50 and inner distance 100: *min* = 160 *max* = 240, read length 100 and inner distance 100: *min* = 260 *max* = 340, read length 100 and inner distance 300: *min* = 380 *max* = 620; *D. simulans* libraries, read length 76: *min* = 176 *max* = 280, read length 120: *min* = 270 *max* = 390). For mrfast we used paired end mapping (-pe), provided a minimum fragment size of 10 (–min), a maximum fragment size of 400 (–max; for the simulated data with a inner distance of 300, −max 700 was used), a maximum number of mismatches of 6 (-e) and required that only the best position of a read should be reported (–best). We specified bam as output format (-b) for ngm and performed a sensitive search (–sensitive; the default is unclear). For novoalign we provided sam as output (-o SAM), set the quality encoding of fastq files to sanger (-o STDFQ), required that a random position is reported for ambiguously mapped reads (-r Random) and provided suitable estimates for the insert size (mean) and the standard deviation of the insert size (sd) (-i mean sd; simulated data: *mean* = 350 *sd* = 50; *D. simulans* libraries, read length 76: *mean* = 228 *sd* = 52, read length 120: *mean* = 396 *sd* = 110). For segemehl we set the maximum insert size to 1500 (-I). For gsnap we used sam as output format (-A sam). Only for the *D. simulans* libraries we set the maximum number of allowed mismatches to 1 (-m 1) as gsnap encountered an error using these data and default settings (Problem sequence; we iteratively removed 5 problem sequences but still encountered the error).

**Table 6:** Overview of the mapping algorithms used in this work. Parameters used for selecting semi-global (g) or local(l) alignments are shown in bold; * see text for more details

|  | Mapper | Version | Parameter | Reference |
| --- | --- | --- | --- | --- |
| global | bowtie2(g) | 2.2.6 | **-end-to-end** -X 1500 | (Langmead and Salzberg, 2012) |
|  | bwa aln | 0.6.2 |  | (Li and Durbin, 2009) |
|  | clc4(g) | 4.4.2.133896 | **-a global** -i -p fb ss *min\**, *max\** | (CLC bio, 2015) |
|  | mrFAST | 2.6.1.0 | -pe -min 10 -max 400* -best -e 6 | (Alkan et al., 2010) |
|  | ngm(g) | 0.4.13 | **-end-to-end** -b -sensitive | (Sedlazeck et al., 2013) |
|  | noalign(g) | 3.03.2 | **-o FullNW** -i *mean\**, *sd\** -F STDFQ -o SAM -r Random | (Novocraft, 2014) |
|  | segemehl | 0.2.0-418 | -I 1500 | (Hoffmann et al., 2009) |
| local | bowtie2(l) | 2.2.6 | **-local** -X 1500 | (Langmead and Salzberg, 2012) |
|  | bwa sw | 0.7.4 |  | (Li and Durbin, 2010) |
|  | bwa mem | 0.7.4 |  | (Li and Durbin, 2009) |
|  | clc4(l) | 4.4.2.133896 | **-a local** -i -p fb ss *min\**, *max\** | (CLC bio, 2015) |
|  | gsnap | 2015-11-20 | -A sam (-m 1)* | (Wu and Nacu, 2010) |
|  | ngm(l) | 0.4.13 | **-local** -b -sensitive | (Sedlazeck et al., 2013) |
|  | noalign(l) | 3.03.2 | -i *mean\**, *sd\** -F STDFQ -o SAM -r Random | (Novocraft, 2014) |

### 3.2 Data sets

We tested the performance of the different alignment algorithms using simulated data and real data.

Simulated paired end data were generated for populations having SNPs with known positions and allele frequencies. This was accomplished in four steps. We first obtained the *D. melanogaster* reference chromosome 2R (r6.03; http://flybase.org/), removed all characters other than A,T,C or G and extracted the first 2Mbp. This small subsequence (the chassis) acted as basis for introducing variants. Second, we generated two modified versions of the chassis: i) we introduced a SNP with a random, not-reference allele all 100bp into the chassis (⇒chassis with SNPs) and ii) we introduced an indel, at a random position with a random Poisson distributed length (*λ* = 1; zero length indels were discarded and Poisson sampling was repeated; insertions had a random sequence), between all pairs of adjacent SNPs into the chassis with SNPs (⇒chassis with SNPs and indels). Third, we generated two sequences serving as templates for simulating paired ends: one consisting of the chassis and the chassis with SNPs (fig. 2A), and another one consisting of the chassis and the chassis with SNPs and indels (fig. 2B). Finally, uniformly distributed paired end reads (equal 5’ distance between consecutive paired ends; uniform base quality of 40) were simulated from these template sequences (fig. 2C). Note that SNPs identified from these data have known positions (each 100bp) and known allele frequencies (ƒ = 0.5). Paired end reads were simulated with SimulaTE (https://sourceforge.net/projects/simulates/; Pandey et al. in preparation) and the number of reads was selected such that a genomic coverage of 200 resulted (*generate-reads paired-end-uniƒormdistribution.py*; ≈ 2 million paired ends for a read length of 100 and 4 million for a read length of 50).

We tested the performance of the different alignment algorithms for real data using paired end reads from a *D. simulans* population that was collected in 2008 in Northern Portugal (Póvoa de Varzim; collected by P. Orozco-terWengel). We established 250 isofemale lines from the population, used one female from each isofemale line and extracted genomic DNA from the pooled flies as described before (Orozco-terWengel et al., 2012). From this DNA we generated two Illumina sequencing libraries. The first was prepared using the Paired-End DNA Sample Preparation Kit (Illumina, San Diego, CA, USA) following fragmentation of the DNA using a nebulizer and size selection using an agarose gel. The library was sequenced on two lanes of an Illumina GAIIx, resulting in 14.3 and 24.7 million 2x76bp paired end reads after trimming [median insert size 232bp; standard deviation of the insert size 25bp; estimated with Picard v1.128 (http://picard.sourceforge.net) after mapping the reads with bwa aln (0.6.2) (Li and Durbin, 2009)].

The second library was prepared with barcoded adapters using a protocol based on the NEBNext®DNA Library Prep Master Mix Set reagents (E6040L) following shearing pooled genomic DNA with a Covaris S2 device (Covaris, Inc. Woburn, MA, USA) and size selection with AMPureXP beads (Beckman Coulter, CA, USA). The library was sequenced on one lane of an Illumina HiSeq 2500 using 2x120bp reads (median insert size 396bp; standard deviation of the insert size 110bp; 84.5 million paired end reads after trimming).

The quality encoding of all reads was converted to Sanger (offset=33) and low quality regions of reads were trimmed with ReadTools (https://github.com/magicDGS/ReadTools −disable-zipped-output −minimum-length 50 −no-5p-trim −quality-threshold 18; per default the quality is converted to Sanger encoding). ReadTools provides a fast implementation of the trimming algorithm described in Kofler et al. (2011a).

We tested whether intersecting of mappers preserves the targets of selection using the data published by Martins et al. (2014). We obtained Illumina paired end data (2 x 100bp) for four populations infected with C-virus for 20 generations (VirSys; accession numbers ERS409784-ERS409787) and for four control populations (ContSys; accession numbers ERS409780-ERS409783).

### 3.3 Data analysis

The simulated reads were mapped to the chassis (see above), the *D. simulans* libraries were mapped to the reference genome of strain M252 (Palmieri et al., 2015) (v1.1; we included the sequences of *Lactobacillus brevis*, *Acetobacter pasteurianus* and two Wolbachia strains; GenBank accession numbers CP000416.1, AP011170.1, AE017196.1, CP001391.1) and the data from Martins et al. (2014) were mapped to the reference genome of *D. melanogaster* (v6.03; we again included the sequences of *Lactobacillus brevis*, *Acetobacter pasteurianus* and two Wolbachia strains). If not mentioned otherwise, mapped reads were filtered for mapping quality (-q 20) and proper pairs (-f 0x002 −F 0x004 −F 0x008; except for the analysis of single end reads) with samtools (v1.2) (Li et al., 2009a). Mapped reads were converted to mpileup files with samtools (v1.2) and the parameters −B −Q 0. SNPs were called using a minimum allele count of 2. The number of true SNPs (every 100th position), the number of false SNPs (not at every 100th position), the frequency of the reference allele (only for true positive SNPs) and the number of extreme outlier SNPs (where the estimated allele frequency deviates by more than 0.4 from the true frequency 0.5) were computed using custom Python scripts (*snp-caller.py*, *stat-snp.py*). For computing allele frequency differences between samples, mpileup files were created with samtools (v1.2; −B −Q 0), the mpileup files were converted to sync files with PoPoolation2 [revision 196; *mpileup2sync.jar* −fastq-type sanger; the minimum quality (–min-qual) was set to 0 for simulated reads and to 20 for *D. simulans* libraries; (Kofler et al., 2011b)], and *F*_ST_ or Fisher exact test p-values (-log10 transformed) were computed with PoPoolation2 (revision 196; ƒ*st-sliding.pl* −min-count 2 −min-coverage 10 −max-coverage 500 −window-size 1 −step-size 1 −suppress-noninformative −pool-size 400 −min-covered-fraction 1.0; fisher-test.pl −min-count 2 −min-coverage 10 −max-coverage 500 −window-size 1 −step-size 1 −min-covered-fraction 1.0). The outlier quantiles of *F*_ST_ and p-values (Fisher exact test; −log10(p-values)] were calculated with Python scripts (ƒ*st-fractionwise.py*).

Differentiation between evolved and control populations for the data from Martins et al. (2014) was assessed with the Cochran-Mantel-Haenszel test (CMH) implemented in PoPoolation2 (Kofler et al., 2011b) (parameters: −min-count 2 −min-coverage 10 −max-coverage 500).

Aligned reads were visually inspected using IGV (Thorvaldsdóttir et al., 2012) and statistical analyses was performed using the R programming language (R Core Team, 2012).

### 3.4 Data availability

The short reads have been made available at the European Nucleotide Archive (ENA; http://www.ebi.ac.uk/ena; PRJEB13602) and the scripts used in this work as well as the simulated reads are available at Dryad (http://datadryad.org/)

## Author’s contributions

RK and CS conceived the study. AML, RK, PN, KAO analysed the data. RK developed the scripts. RK, AML and CS wrote the paper.

## Acknowledgements

We thank all members of the Institute of Population Genetics for feedback and support. This work was supported by the European Research Council Grant “ArchAdapt” and Austrian Science Funds (FWF-W1225).

